# FACTORs: a Novel R Package for Functional Data Reduction and Hypothesis Testing

**DOI:** 10.1101/2022.10.28.514171

**Authors:** John S. Malamon, Jessica L. Saben, Bruce Kaplan

**Affiliations:** University of Colorado Anschutz Medical Campus, Department of Surgery; Colorado Center for Transplantation Care, Research and Education (CCTCARE); University of Colorado Anschutz Medical Campus, Department of Medicine

## Abstract

To extract biological meaning from transcriptomics analysis, investigators almost exclusively rely on biological annotation databases and the subsequent associations made between gene products and ontology terms (i.e., gene ontology analysis). Although there is ease and utility to this approach, multiple hypothesis testing methods such as differential expression analysis are performed in the absence of *in silico* validation, and downstream gene ontology analysis methods lack precision and propagate Type I error. Therefore, we present a novel R package, Functional Association Vectors (FACTORs), that begins to address some of the common pitfalls associated with functional association studies in transcriptomics research. FACTORs are vectorized containers for directly comparing and testing congruent functional association statistics at the molecular level. Our goal was to develop novel methodology an R software package to allow for the experimental validation of differentially expressed genes that are conserved across studies and to reduce Type I error in the down-stream functional analysis of these signals. FACTORs are generalizable and flexible, allowing for any association statistic such as log fold change (logFC), t-statistic, p-value, or other functional significance score. To demonstrate utility of FACTORs in the cross-study validation of global mRNA expression profiling, we used differential expression analysis summary statistics obtained from two studies with publicly available transcriptomic data. Through this demonstration we show FACTORs provide a more precise and generalizable functional hypothesis testing methodology and data reduction approach that directly tests functional association statistics at the molecular level, across experiments.

**AUTHOR SUMMARY:** We present a novel statistical methodology and software tool that can be used to validate transcriptomic data across studies. FACTORs is available as an R package and will aid in transcriptomic data reduction, identifying gene expression profiles that are conserved across studies, and improving the precision and generalizability of functional association studies. We propose FACTORs as a generalizable methodology for reducing Type I error and increasing the biological relevance of functional associations in transcriptomics studies.

## INTRODUCTION

In assigning biological functions to experimentally derived gene sets and their products such as mRNA transcripts, genes, and proteins, investigators almost exclusively rely on biological annotation databases and the subsequent associations made between gene products and ontology terms. This is referred to as gene ontology analysis and involves querying against massive biological databases consisting of hundreds of millions of annotation terms (nodes). Examples of such databases are the Gene Ontology consortium (GO)[1] and Kyoto Encyclopedia of Genes and Genomes (KEGG)[2]. Enrichment or over-representation analyses, which compare the observed number of gene products to the expected in all possible annotation graphs[3], are commonly used to generate associations between experimentally derived gene products and ontology terms. With this methodology, one can directly test the gene expression changes in relation to biological states and functions, reducing false-positive associations.

Although there is ease and utility to this approach, there are many pitfalls that should not be overlooked[4, 5]. Importantly, gene ontology analysis methods lack precision and propagate Type I error. First, the defining background characteristics and distributions of the data sources being conflated to derive these complex annotation graphs are not maintained. Typically, investigators employ various statistical methods to control known heterogeneities and batch effects based on the descriptive characteristics of background data and sample variation to account for such differences. However, this is not possible with large-scale annotation databases because study and sample-level details are not retained. While transitivity is assumed, these methods often produce results that are out of context regarding the biological mechanisms and experimental conditions under study, leaving investigators with the difficult decision of what results to include in their publication. Second, gene ontology analysis often produces a broad spectrum of highly significant, but still unspecific results that are cumbersome and time-consuming to interpret. This very often leads to inflation bias or “p-hacking”, where investigators run gene ontology analysis using various databases and only keep statistically significant results that fit the expected outcome. This also introduces confirmation bias. Other issues that plague gene ontology analysis are inter-ontology links and annotation redundancies, which increase Type I error.

Here we present FACTORs (<u>F</u>unctional <u>A</u>ssociation Vec<u>tors</u>), a novel R package for cross-study validation, functional data reduction, and hypothesis testing. FACTORs provide a more precise and generalizable functional hypothesis testing methodology and data reduction approach that directly tests functional association statistics at the gene product level, across experiments. We propose FACTORs R package as a new paradigm for reducing Type 1 error and increasing the biological relevance of functional associations in an era where multiple hypothesis testing and Type I error are largely uncontrolled.

## RESULTS

### Use-Case Overview

Very often, multiple hypothesis testing methods such as differential expression analysis are performed in the absence of *in silico* validation. Moreover, it is extremely useful to reduce the hundreds and thousands of statistically significant results obtained using differential expression analysis to a manageable set of hypotheses that can be validated using orthogonal methods. Our goal was to develop an R package that will: 1) efficiently validate differentially expressed genes that are conserved across multiple studies and 2) reduce Type I error in the down-stream functional analysis of these signals. **Figure 1** provides a general description of the analytical workflow that can be used with the FACTORs R package to perform cross-study validation. One of the major advantages to the FACTORs R package is that multiple data input types can be used (i.e., raw sequence reads, microarray data, or summary statistics). ‘Example study 1’ illustrates the workflow that would be used for raw mRNA sequence data, whereas ‘Example study 2’ provides the workflow that would be used if starting with summary statistics from a differential expression analysis. FACTORs R package provides the R code necessary to take each of these types of input data to a position where validation across studies can be performed. Once a list of differentially expressed genes has been created for each study (depicted in the red and green boxed-in tables, **Figure 1**), FACTORs are constructed, and CCF hypothesis testing is deployed. These novel methods work to reduce the target gene list to a group of differentially expressed genes that are conserved in each study. With the resultant list of overlapping genes, gene ontology analysis can be utilized (e.g., GO or KEGG) with greater confidence and specificity. Thus, FACTORs R package provides a simple paradigm for functional hypothesis testing and Type 1 error reduction by directly evaluating functional association statistics across studies.

**Figure 1.**
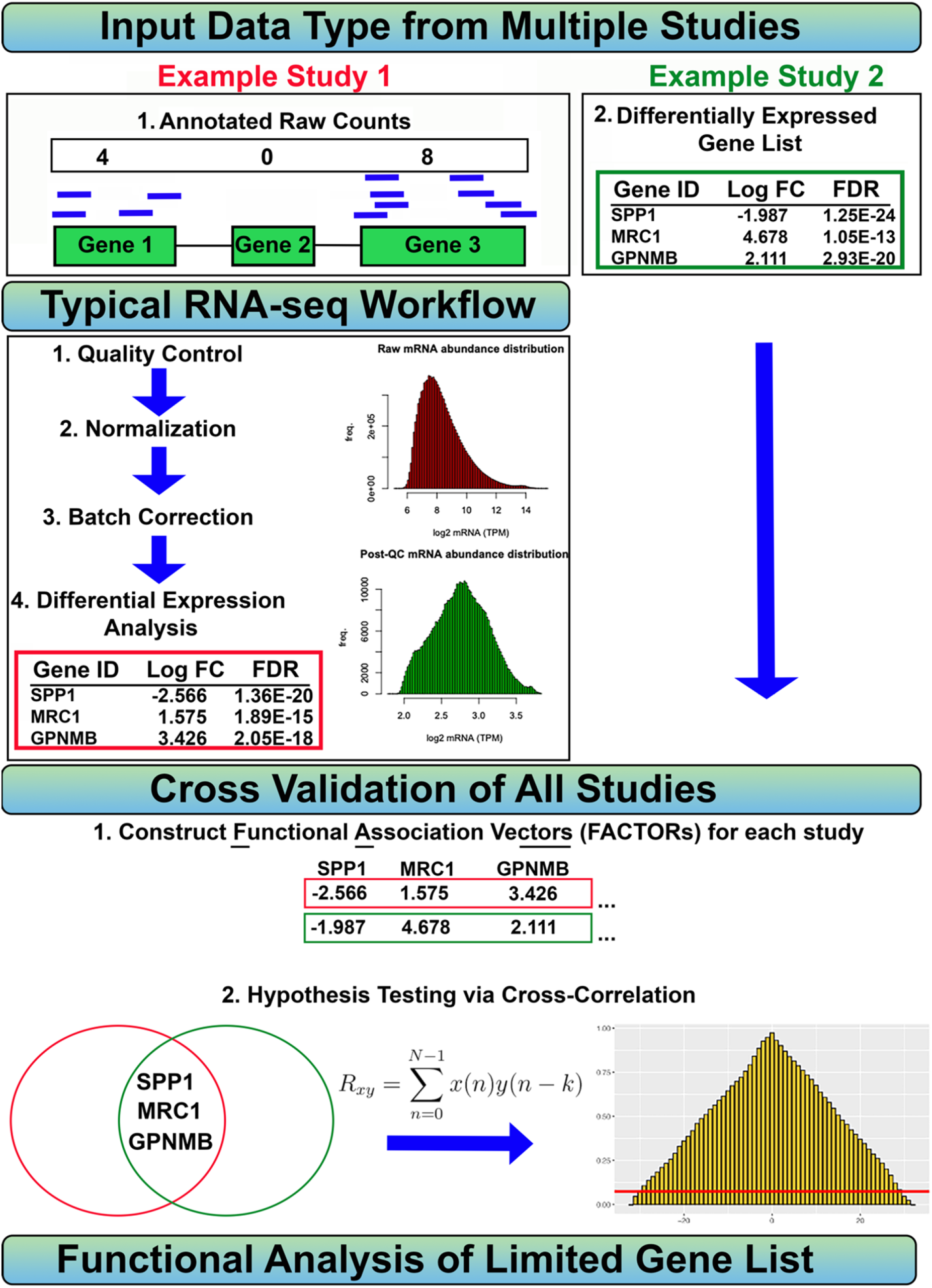
Overview of Functional Association Vectors or FACTORs framework. Our novel framework consists of five main analysis components: quality control, normalization, batch correction, differential expression analysis, and functional cross-correlation hypothesis testing. FACTORs are the ordered functional association statistics of experimental transcriptional datasets and can be constructed from summary statistics supplied via functional association, rank, or weighting scores. FACTORs can also be constructed from high-dimensional scRNA counts using differential expression analysis. FACTORs provide a precise, standardized, and flexible methodology for defining and directly testing functional associations, terms, and ontologies at the molecular level across experiments.

### Demonstration of Cross-study Validation using FACTORs and the CCF Hypothesis Testing

To demonstrate the ease and utility of our approach in the cross-study validation of global mRNA expression profiling, we constructed FACTORs and performed CCF hypothesis testing on differential expression analysis summary statistics obtained from two studies. First, we obtained raw microarray data provided by Abbas *et al*. or ‘Example Study 1’[6] constructed from mRNA expression profiles of 22 PBMC types. Using FACTORs R package, we performed QC, normalization, and batch correction on these data (GS22886). To examine conserved markers specific to T cells that are positive for cluster of differentiation 8 (CD8+) expression, we trained CD8+ T cells against 21 other immune cell types in the differential expression analysis [6] and obtained 559 unique differentially expressed genes with a p-value less than 0.001 (**Supplemental_Spreadsheet_01.xlsx**). To cross validate these findings, we then downloaded the CD8+ T cell marker gene summary statistics provided by Koutsakos et. al [7] via the SCEA database[8, 9]. These summary statistics (‘Example Study 2’, **Figure 2**) contained 149 unique CD8+ T cell transcripts with a p-value less than 0.001 (**Supplemental_Spreadsheet_02.xlsx**). Next, we used enrichR to perform a gene ontology analysis using the Human Gene Atlas on the resulting gene sets from each differential expression analysis. ‘Example Study 1 yielded 117 associated genes overlapping the ‘CD8+ T cells’ ontology term with an odds ratio of 10.35 and adjusted p-value of 1.5e-63, whereas gene ontology analysis using summary statistics (Example Study 2) yielded 20 associated genes with an odds ratio of 5.13 and adjusted p-value of 8.7e-7 (**Figure 2B**). Next, FACTORs were constructed from the two example gene sets. These two sets of differentially expressed genes (Example Study 1 and Example Study 2) shared 17 out of 692 distinct genes (**Figure 2**). FACTORs analysis provided a hypergeometric p-value (N=20,000) of 4.0e-32. We then performed the same gene ontology analysis on the 17 shared CD8+ marker genes and found that the ‘CD8+ T cells’ ontology had 10 associated genes, an odds ratio of 46.79, and an adjusted p-value of 1.1e-10. FACTORs provided a 4.52 and 9.12-fold increase in the functional association odds ratio for the ‘CD8+ T cells’ ontology, respectively. Finally, we performed CCF hypothesis testing (**Eq. 5 and Figure 2C**) on these two FACTORs to derive a CCF p-value of 0.0001. The CCF significance threshold was 0.074 (**Eq. 6 and Figure 2C**). This use-case demonstration shows that by employing the FACTORs R package we were able to identify and validate a smaller gene set list associated with CD8+ T cells from two completely different studies, providing increased confidence in the identified genes as conserved markers of CD8+ T cells.

**Figure 2.**
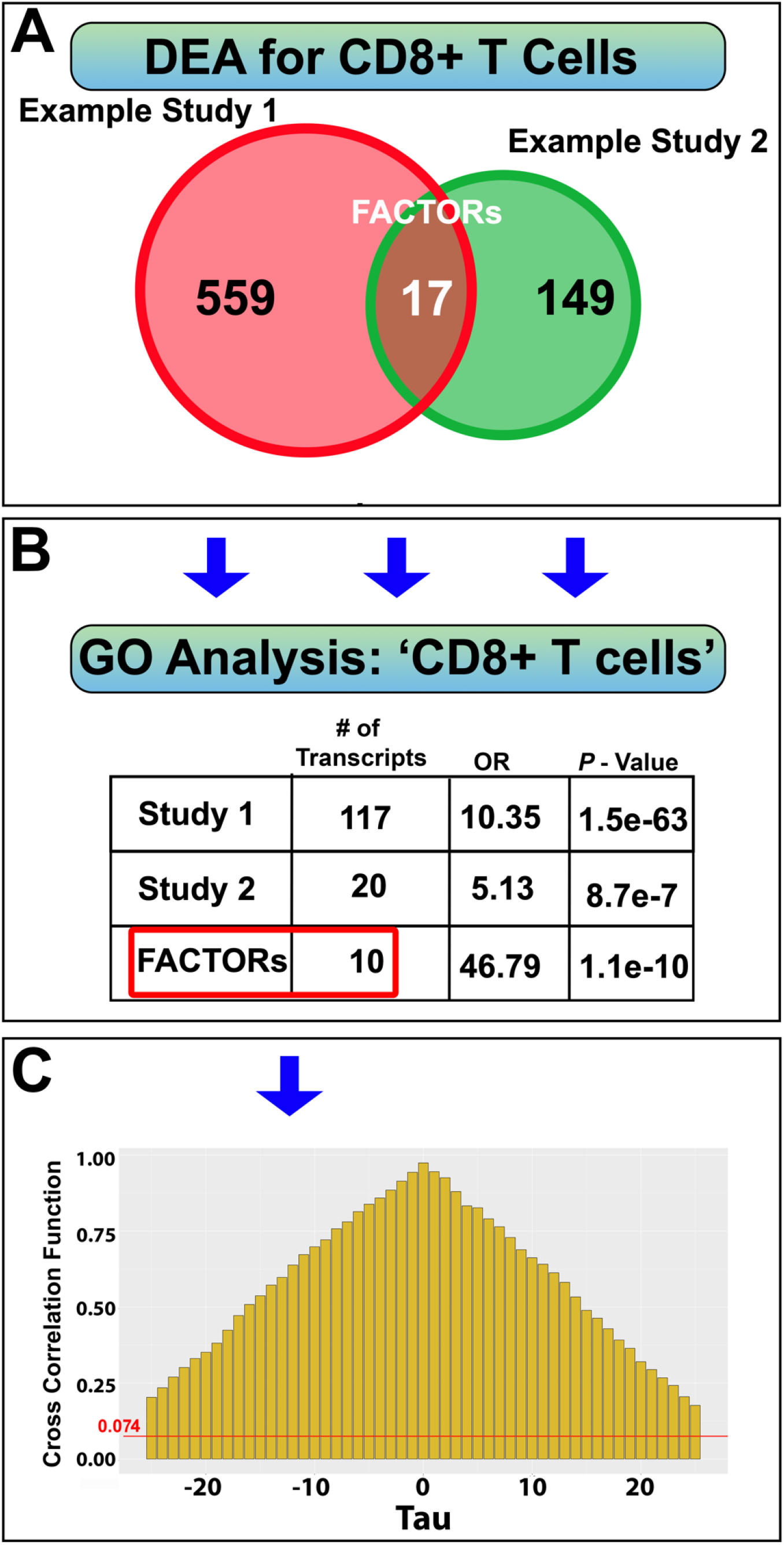
Demonstration of Cross-study Validation using FACTORs and CCF Hypothesis Testing. To demonstrate our approach in the cross-study validation of global mRNA expression profiling, we constructed FACTORs and performed CCF hypothesis testing on differential expression analysis summary statistics obtained from two sources, GS22886 and the SCEA database. We obtained 559 and 149 unique differentially expressed genes with a p-value less than 0.001, respectively. FACTORs were constructed from the two DE gene sets. These two sets of DE genes shared 17 out of 692 distinct genes, providing a hypergeometric p-value (N=20,000) of 4.0e-32. We then performed the same gene ontology analysis on the 17 shared CD8+ marker genes and found that the ‘CD8+ T cells’ ontology had 10 associated genes, an odds ratio of 46.79, and an adjusted p-value of 1.1e-10. FACTORs provided a 4.52 and 9.12-fold increase in the functional association odds ratio for the ‘CD8+ T cells’ ontology, respectively. Finally, we performed CCF hypothesis testing on these two FACTORs to derive a CCF p-value of 0.0001. The CCF significance threshold was 0.074

## MATERIALS AND METHODS

### Data sources

This study utilized data from two publicly available sources. The first is Abbas *et al*. (2005)[6], which provided microarray raw data and summary statistics for twelve distinct human leukocyte cell types derived from peripheral blood mononuclear cells (PBMCs). These data are available via the Gene Expression Omnibus (GEO)[10] under accession GSE22886. All cells were analyzed using the Affymetrix Human Genome U133A/B array. The main objective of this study was to provide a high-quality transcriptomic dataset to decipher gene expression patterns in resting and activated immune cells. The second data source used in this study was obtained via the Single Cell Expression Atlas (SCEA)[8, 9] as published by Koutsakos *et al*. [7]. The SCEA is a comprehensive, single-cell gene expression database provided by the European Molecular Biology Laboratory’s European Bioinformatics Institute. Currently, the SCEA contains single-cell RNA sequencing for twenty species and over 300 studies and 8.5 million unique cell lines.

### Statistical approach

#### Hypergeometric Probably and Enrichment Factor

Two sets of gene products sets are generated and have *n* overlaps. We want to ask, what is the probability of observing *n* overlapping genes in two independent sets without replacement? *Eq. 1* provides the hypergeometric probability function, where *n* is the number of overlapping genes, *GS_1_* and *GS_2_* are two independent gene sets, and *N* is the entire gene pool. The enrichment factor is the number of matching genes (*n*) divided by the expected number of overlapping genes if randomly selected. An enrichment factor greater than one indicates more overlap than expected at random. The enrichment factor is commonly used in gene ontology analysis.

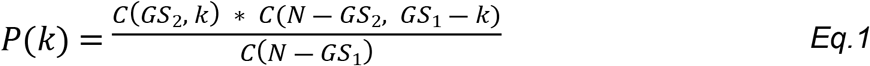

#### FACTORs: Containers for the Cross-study Validation of Functional Association Statistics

FACTORs are vectorized containers for directly comparing and testing functional association statistics (*a_j_*), where x_j_ represents a unique gene product. FACTORs are generalizable and flexible, allowing for any association statistic such as log fold change (logFC), t-statistic, p-value, or other functional significance score. Functional significance could also be defined by variable weighing methods such as the Random Forest method or other tree-based models. Treating FACTORs as sets as congruent association statistics is a natural extension of many NGS analysis frameworks and standardizes functional definitions and their applications. FACTORs integrate seamlessly with existing tools and analysis methods to provide precise functional definitions for in- and cross-study analysis and validation.

#### Use Cases for Data Reduction and Hypothesis Testing

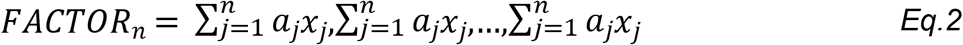

Where, *FACTOR_n_* is a vector of functional association statistics (*a_j_*) for all genes (x_j_) provided by a functional analysis. P-value (x_j_) vectors can be transformed to t-statistics using *Eq.3*.

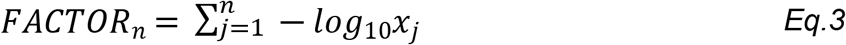

#### Data Reduction via Cross-Correlation of Experimentally Derived Functional Association Statistics

FACTORs are ordered functional association statistics of gene sets. For any two FACTORs, we perform hypothesis testing by applying the discrete cross-correlation function (CCF) provided in *Eq. 4*. Non-overlapping genes are filtered out. When applying the CCF, we recommended constructing FACTORs using the same association methods.

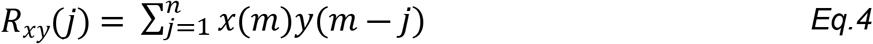

#### Calculating the Cross-Correlation Functional P-value

*Eq. 5* provides the two-sided p-value of the CCF (*R_xy_*), under the null hypothesis and the assumption of normality, where the mean of the CCF is 0 and its standard deviation is equal to 1/*N*. Here, *N* is equal to the number of shared observations between two FACTORs. The CCF p-value is the probability that the max correlation coefficient is greater than the maximum observed correlation provided in *Eq. 4*.

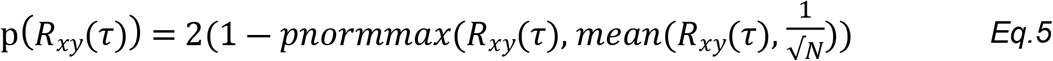

Where, *p(R_xy_(τ))* is the two-sided probability that the null hypothesis (CCF=0) is greater than the observed cross-correlation coefficients. Finally, the CCF significance threshold is provided in *Eq. 6*. A maximum cross-correlation coefficient greater than the CCF significance threshold implies that we reject the null hypothesis that two sets of functional association statistics are not correlated above chance. Furthermore, we can scale the CCF significance threshold up to be more selective by increasing the denominator of *Eq.6*.

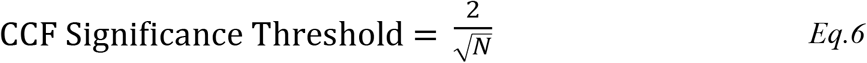

## DISCUSSION

Transcriptomics is a powerful tool used to understand differences in cell types or varying biological paradigms. However, the amount of data generated from these analyses can be daunting for some investigators, which can then become magnified by the numerous highly significant, yet largely unspecific findings. Gene ontology analysis is the gold standard method used to extrapolate experimentally derived gene expression changes to potential biological functions. However, several statistical pitfalls, such as increased potential for Type 1 error and inflation bias are associated with these downstream analyses. Herein, we present a novel R package, FACTORs, that can begin to address some of the common pitfalls associated with functional association studies and will increase the validity and reproducibility of global mRNA expression profiling.

Transcriptomics has been used by molecular biologists across scientific disciplines since the commercialization of the microarray in the late 1990s[11]. More recently, global mRNA expression studies have adopted RNA-sequencing strategies as sequencing technologies have become both cheaper and more readily available[12]. Interestingly, the transcriptomic field has not developed a consensus on how to validate large mRNA expression datasets. Historically, real-time quantitative polymerase chain reaction (RT-qPCR) was used to internally validate microarray data using the same cDNA samples that were applied to the microarray[13]. However, the utility of this approach is not always clear, especially with new transcriptomic strategies such as RNA-sequencing[14]. Alternatively, orthogonal approaches such as RT-qPCR may be useful for confirming differential gene expression across studies and/or within differing experimental paradigms. However, these approaches are not always possible or applicable to the experimental scenario. While several other ‘omics’ fields have implemented guidelines for including validation cohorts as a prerequisite for publication[15], the field of transcriptomics has yet to adopt this practice. The FACTORs R package presents a novel way to do cross-study validation *in silico* making the option to validate findings cost effective and feasible. To improve on the reproducibility and transparency of “big data” research, may journals require data sharing on public platforms[16] such as GEO and SCEA. The call for data sharing in omics research began over a decade ago[15, 17] and therefore these databases contain a rich source of transcriptomics data that can, and should, be used for cross-study validation.

As with all methodologies, this approach has limitations and must be evaluated within the context of the data used in this study. First, we tested this methodology with transcript levels derived from microarray platforms. We could not obtain replication studies sequenced using RNA-seq; however, these data will become available. Another limitation of this study is the sample size. In this study, we were limited to no more than a dozen sample for each cell type or state. Again, we will apply this methodology to additional sample as data become more abundant. Finally, this methodology require some limited programming skills and is more complex than GOA.

In conclusion, the FACTORs R package presented herein provides a novel, user-friendly, statistical method for performing cross-validation of transcriptomics studies *in silico*. FACTORs R package can resourcefully validate transcriptional profiles that are conserved across multiple studies, reducing differentially expressed gene lists to more reasonable and confident gene sets that have significantly improved functional association odds ratios in gene ontology analysis. Finally, the FACTORs theory and associated R code has potential usage beyond mRNA analysis and can be adapted to other big data applications.

## ACKNOWLEDGEMENTS

N/A

## DATA AVAILABILITY STATEMENT

The FACTORs R package can be found at the following link: https://github.com/jmal0403/FACTORs. A detailed wiki with code example can be found here: https://github.com/jmal0403/FACTORs/wiki.

## ABBREVIATIONS

CCF: Cross-Correlation Function
CD8: Cluster of Differentiation 8
FACTORs: Functional Association Vectors
GEO: Gene Expression Omnibus
GO: Gene Ontology
KEGG: Kyoto Encyclopedia of Genes and Genomes
PBMC: Peripheral Blood Mononuclear Cells
RT-qPCR: Real-Time Quantitative Polymerase Chain Reaction
SCEA: Single Cell Expression Atlas

